# Analyses of spike protein from first deposited sequences of SARS-CoV2 from West Bengal, India

**DOI:** 10.1101/2020.04.28.066985

**Authors:** Feroza Begum, Debica Mukherjee, Dluya Thagriki, Sandeepan Das, Prem Prakash Tripathi, Arup Kumar Banerjee, Upasana Ray

## Abstract

India has recently started sequencing SARS-CoV2 genome from clinical isolates. Currently only few sequences are available from three states in India. Kerala was the first state to deposit complete sequence from two isolates followed by one from Gujarat. On April 27, 2020, the first five sequences from the state of West Bengal (Eastern India) were deposited on ‘a global initiative on sharing avian flu data’ (GISAID) platform. In this paper we have analysed the spike protein sequences from all these five isolates and also compared for their similarities or differences with other sequences reported in India and with isolates of Wuhan origin. We report one unique mutation at position 723 and the other at 1124 in the S2 domain of spike protein of the isolates from West Bengal only and one mutation downstream of the receptor binding domain at position 614 in S1 domain which was common with the sequence from Gujarat (a state of western part of India). Mutation in the S2 domain showed changes in the secondary structure of the spike protein at region of mutation. We also studied molecular dynamics using normal mode analyses and found that this mutation decreases the flexibility of S2 domain. Since both S1 and S2 are important in receptor binding followed by entry in the host cells, such mutations may define the affinity or avidity of receptor binding.

## Introduction

SARS-CoV2 (a member of Coronaviruses) outbreak occurred in Wuhan, China in the year 2019 and it became a pandemic recently that has affected countries worldwide. To design antiviral therapeutics/ vaccines it is important to understand the genetic sequence, structure and function of the viral proteins. When a virus tries to adapt to a new environment, in a new host, in a new geographical location and a new population, it would make changes in its genetic make up which in turn would bring in slight modifications in the viral proteins. Such variations would help the virus to utilize the host’s machinery the best in favour of the virus survival and propagation. Since host’s immune system eventually learns to identify a pathogen that had infected and starts producing protective antibodies, a virus often changes its structural proteins such that it can still infect the host cells escaping the host’s immune system. Coronaviruses have been long known to undergo rapid mutations in its RNA genome. Such mutations get reflected in changes in the amino acid sequences of its structural and the non-structural proteins.

Spike protein is one of the structural proteins of SARS-CoV2 that forms a homotrimer on the surface of the vital lipid envelope. This trimer is made up of monomers consisting of S1 and S2 subunits. S1 subunit helps in attachment to the host cell receptor while S2 subunit helps in fusion to host cell and entry. Thus, spike protein has been an area of interest for designing vaccine and antiviral candidates against SARS-CoV2. Since the spike protein tends to mutate, it is important to obtain a broad mutation profile of this protein from extensive genome sequencing from different geographical locations of the world. Targeting areas of spike protein that do not undergo mutation i.e. conserved would be the key to design effective broad-spectrum antiviral or vaccine.

## Methods

### Sequences

We downloaded the five new SARS-CoV2 sequences from West Bengal (EPI_ISL_430468; EPI_ISL_430467; EPI_ISL_430465; EPI_ISL_430464; EPI_ISL_430466) from GISAID database and the spike protein sequences corresponding to Kerala isolates [2] and Gujarat isolate [3] from the NCBI virus database.

For the West Bengal isolates, the nucleotide sequences corresponding to the spike protein were selected and then translated on ‘ExPASY’ Translate tool to obtain the protein sequences.

### Sequence alignments and structure

All the spike protein sequences were aligned by multiple sequence alignment platform of CLUSTAL Omega. The alignment file was viewed using MView and differences in the sequence or the amino acid changes were recorded.

CFSSP (Chou and Fasman secondary structure prediction) server was used to predict secondary structures of SARS-CoV2 spike protein.

To study the effect of mutation on the conformation, stability and flexibility of the spike protein, structure was downloaded from RCSB PDB. We used the available SARS-CoV-2 spike ectodomain structure (open state) (PDB ID: 6VYB). 6VYB structure was uploaded on DynaMut software (University of Melbourne, Australia) [4] and change in vibrational entropy; the atomic fluctuations and deformation energies due to mutation were determined. For atomic fluctuation and deformation energy calculations, calculations were performed over first ten non-trivial modes of the molecule.

## Results and Discussion

The first set of five sequencing data from clinical isolates of SARS-CoV2 from the state of West Bengal, India was submitted on 27.2.2020 by National Institute of Biomedical Genomics (NIBMG) in collaboration with ICMR-National Institute of Cholera and Enteric Diseases (ICMR-NICED). The sequences were submitted on GISAID database.

We downloaded all the sequences from West Bengal (Figure 1) and performed a nucleotide translation to obtain respective spike protein sequences. All these spike protein sequences were first aligned in CLUSTAL Omega to check for similarities or differences. We found that all the isolates from West Bengal were identical (data not shown as they were identical). So, we used one of these sequences as representative of SARS-CoV2 spike from West Bengal for our further analyses.

**Figure 1:**
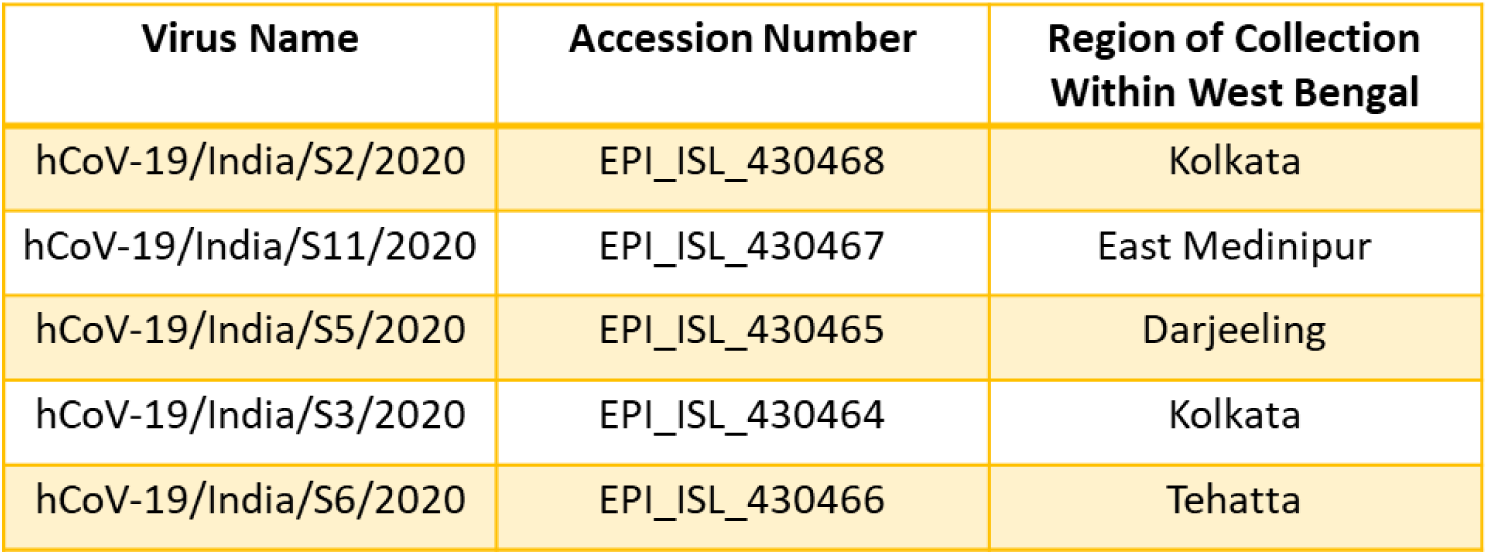
Table represents tabulation of accession numbers of the sequences from kolkata used for the anlayses with specific regions of smaple collection

Since, currently we have sequences against SARS-CoV2 only from three states in India i.e. Kerala, Gujarat and now West Bengal, we compared all the sequences to detect possible changes (Figure 2). We considered the original Wuhan sequence as the wild type for comparison. Based on these criteria we found four different amino acid positions that were mutated in these isolates overall. We had recently published the details with respect to the spike protein mutations in Kerala and Gujarat isolates.

**Figure 2:**
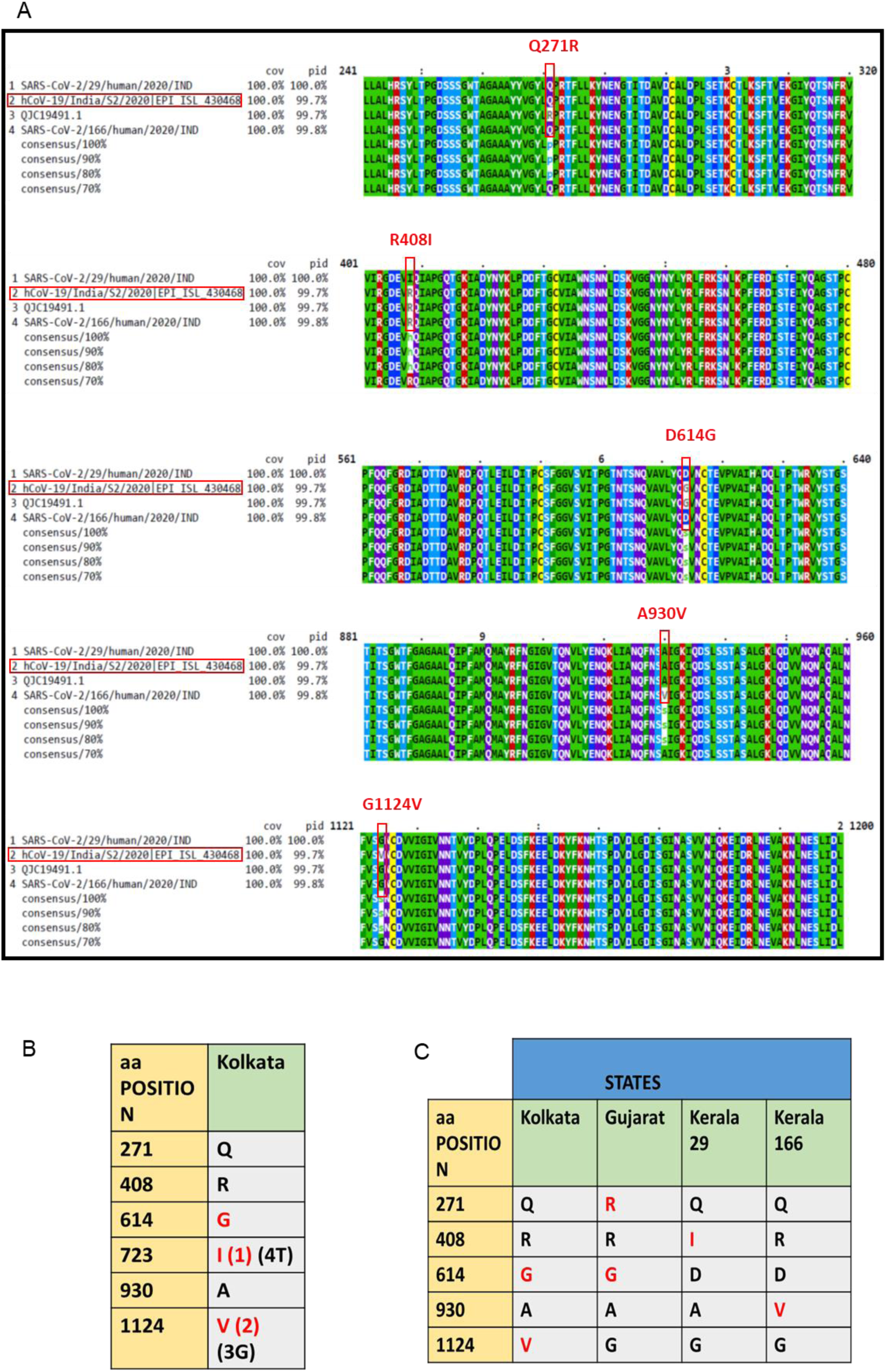
Mutation analysis of isolates from Kolkata, Gujarat and Kerala. A. Multiple sequence alignment of Spike protein sequence of Kolkata isolate with sequences obtained from other parts of India. Sites of mutation are showed in Red. B. Tabulation of amino acid mutations among isolates form Kolkata. Mutations are showed in red. Number/s in parenthesis shoe number of isloates that showed the aminoacid type. C. Tabulation of amino acid at the points of mutation for isolates from different parts of India.

Here, we report that among West Bengal isolates there were three mutations in the spike protein. One of these mutations was D614G in the S1 domain. This lies near the receptor bending domain at a downstream position. The other mutation was G1124V in the S2 domain. While D614G was also found in the isolate from Gujarat but not in Kerala isolates, G1124V mutation was exclusively found in two of the isolates of West Bengal. None of the isolates from other parts of India had this mutation. Another mutation that appeared in one of the West Bengal’s isolates was T723I.

We characterized mutation G1124V.Both glycine (G) and valine (V) are non-polar amino acids with aliphatic R groups. Glycine has no side chain whereas valine is bulkier due to its side chain. A change from glycine to valine can thus potentially disrupt the local folding of the protein. For example, it was shown that G to V change in a P-glycoprotein changed its drug specificities [5].

Secondary structure prediction showed changes in and around the site of mutation (Figure 3). In the mutant spike there was a loss of turn structure from position 1124 and addition of four helices at positions 1123, 1124, 1125 and 1126. This change in secondary structure might lead to change in function of S2. S2 helps in fusion process of the spike protein and thus mutation in S2 may have altered receptor spike interactions and thus infectivity.

**Figure 3:**
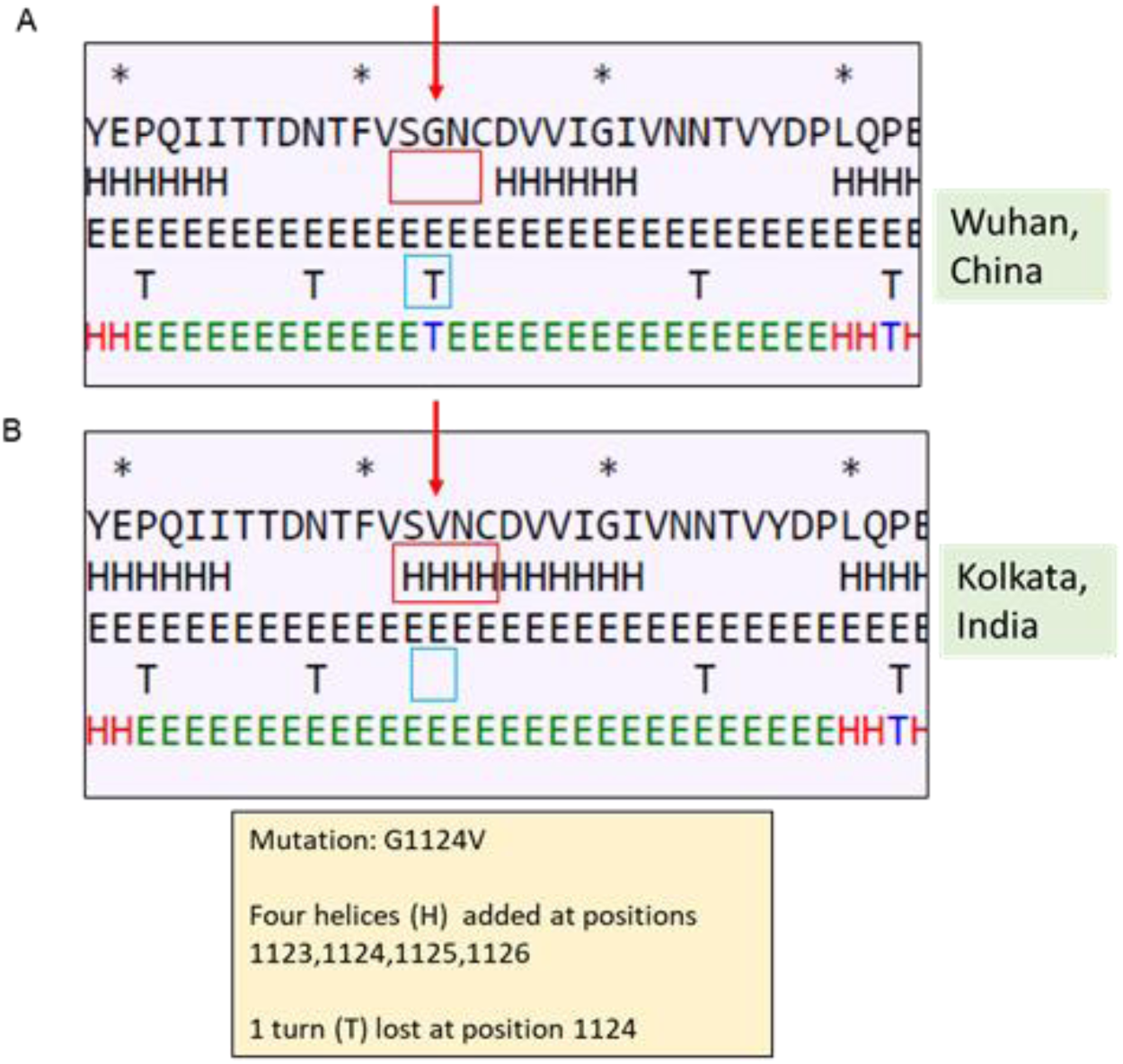
Effect of mutation at position 1124 on secondary structure of Spike protein. A. Secondary structure of Spike protein of Wuhan isolate (the area around the residue 1124 has been shown) B. Secondary structure of Spike protein of Kolkata isolate shoeing effect of mutation on the secondary structure.

To correlate if changes in secondary structure also gets reflected in the dynamics of the protein in its tertiary structure, we performed normal mode analyses and studies protein stability and flexibility. Change in vibrational entropy energy (ΔΔSVib ENCoM) between the wild type Wuhan isolate and the West Bengal isolate was −4.445 kcal.mol^−1^.K^−1^(Figure 4). The ΔΔG was 0.905 kcal/mol and the ΔΔG ENCoM was 4.756 kcal/mol. All these suggested a stabilizing mutation in this type of spike. The interatomic interactions have been shown in Figure 5. Analyses of atomic fluctuations and deformation energies showed visible changes (Figure 6). Atomic fluctuations calculate the measure of absolute atomic motion whereas the deformation energies detect the measure of flexibility of a protein. Figure 6 shows the visual representations of the atomic fluctuation and deformation energies where positions that could be visibly detected to be different have been marked.

**Figure 4:**
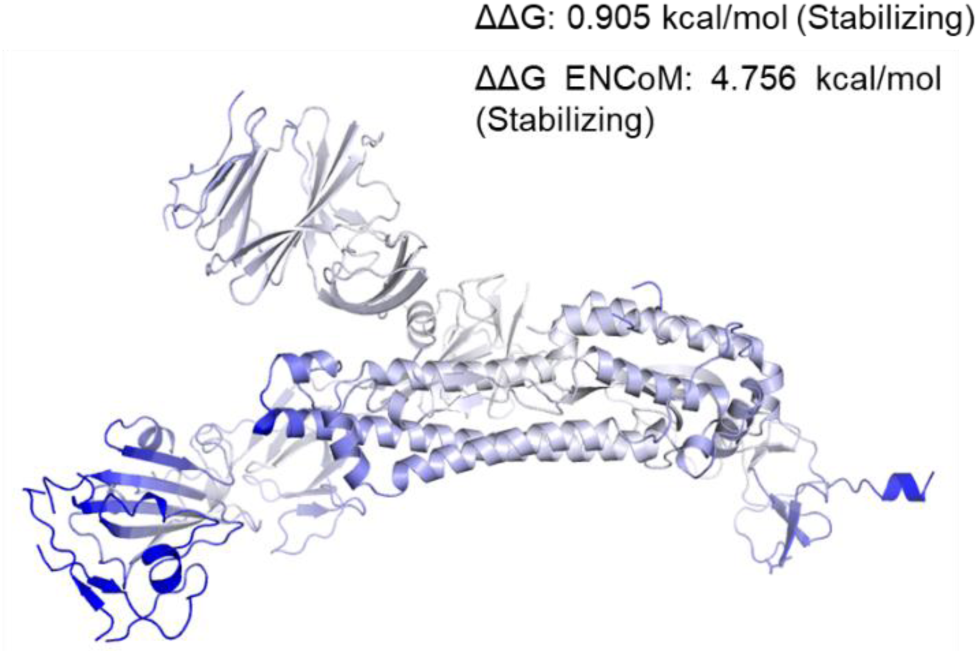
Δ Vibrational Entropy Energy Between Wild-Type and Mutant. Δ Vibrational Entropy Energy Between Wild-Type and Mutant ΔΔS_Vib_ ENCoM: −4.445 kcal.mol^−1^.K^−1^. Amino acids were coloured as per the vibrational entropy change due to mutation. Blue represents rigidification of structue

**Fig. 5:**
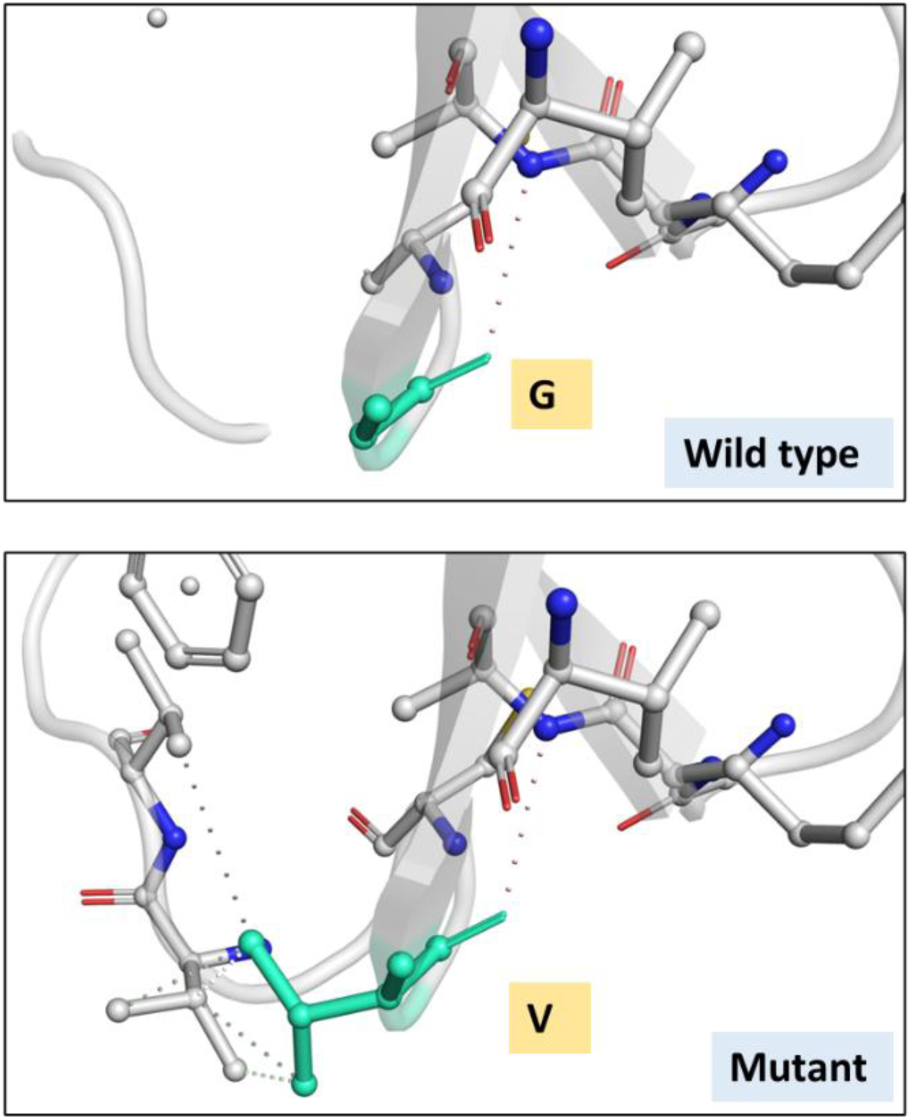
Interatomic Interaction. Wild-type and mutant residues are colored in light-green and are represented as sticks alongside with the surrounding residues which are involved on anytype of interactions

**Fig. 6:**
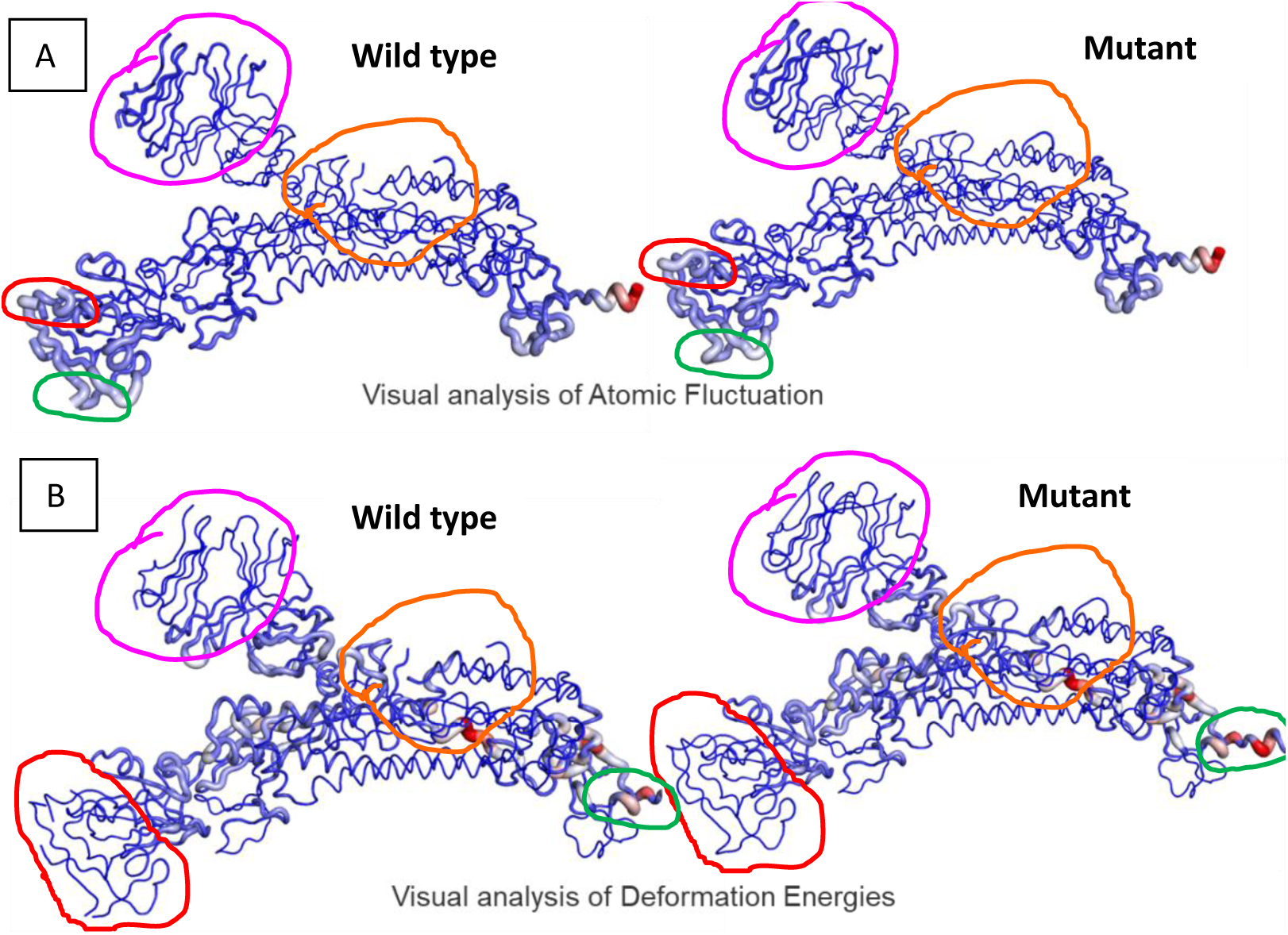
A. Analysis of atomic fluctuations B. Visual analysis of deformation energies. Magnitude of fluctuation and deformation has been shown using thin to thick tube coloured blue (low), white (moderate) and red (high)

This is the first report of mutations of such types in the isolates of the state of West Bengal and further sequencing followed by sequence analyses would help expanding the knowledge about variations of spike protein in human SARS-CoV. These variations might lead to virus diversification and eventual emergence of variants/ antibody escape mutants/strains/ serotypes. Also, mutations might help the virus to expand its tissue tropism and adjust with the host environment better. Therefore, elaborate studies on sequence variations should be done which would in turn help in better therapeutic targeting.

## Acknowledgements

We thank Prof. Saumitra Das, NIBMG, Kalyani, West Bengal and ICMR-NICED for depositing the sequences in public database and making it available open access. We thank CSIR, AcSIR and North Bengal Medical College and Hospital for other necessary support and input.

